# SYNAPTIC DYSFUNCTION UNDERLIES ALTERATIONS IN THE INITIATION OF GOAL-DIRECTED BEHAVIORS: IMPLICATIONS FOR HIV-1 ASSOCIATED APATHY

**DOI:** 10.1101/2021.09.02.458761

**Authors:** Kristen A. McLaurin, Michael N. Cranston, Hailong Li, Charles F. Mactutus, Steven B. Harrod, Rosemarie M. Booze

## Abstract

Individuals living with human immunodeficiency virus type 1 (HIV-1) exhibit an increased prevalence of neuropsychiatric comorbities (e.g., apathy) relative to their seronegative counterparts. Given the profound functional consequences associated with apathy, conceptualizing the multidimensional neuropsychiatric syndrome, and associated neural mechanisms, following chronic HIV-1 viral protein exposure remains a critical need. HIV-1 associated apathy was examined by quantifying goal-directed behaviors, indexed using voluntary wheel running, during the diurnal and nocturnal cycle. Apathetic behaviors in the HIV-1 Tg rat were characterized by a profound decrease in the number of running bouts during both the diurnal and nocturnal cycle, supporting a prominent deficit in the self-initiation of spontaneous behaviors. Additionally, HIV-1 Tg animals exhibited a decreased reinforcing efficacy of voluntary wheel running during the nocturnal cycle. Following the completion of voluntary wheel running, synaptic dysfunction in medium spiny neurons (MSNs) of the nucleus accumbens (NAc) was examined as a potential neural mechanism underlying HIV-1 associated apathy. HIV-1 Tg animals displayed prominent synaptic dysfunction in MSNs of the NAc, characterized by decreased synaptic connectivity and a population shift towards an immature dendritic spine phenotype relative to control animals. Synaptic dysfunction accounted for 42.0% to 68.5% of the variance in the number of running bouts affording a key neural mechanism underlying the self-initiation of spontaneous behaviors. The establishment of a fundamental neural mechanism underlying apathy affords a key target for the development of novel therapeutics and cure strategies for affective alterations associated with HIV-1.

## INTRODUCTION

Despite widespread implementation of combination antiretroviral therapy (cART), the primary treatment regimen for individuals with human immunodeficiency virus type 1 (HIV-1), neurocognitive (i.e., HIV-1 associated neurocognitive disorders (HAND); for review, Eggers et al., 2017) and neuropsychiatric (e.g., depression, apathy; for review, Cysique & Brew, 2019) comorbidities persist. Apathy, specifically, afflicts between 26-42% of HIV-1 seropositive individuals in the post-cART era (Tate et al., 2003; Kamat et al., 2012); prevalence rates which are significantly greater than their seronegative counterparts (Kamat et al., 2012; Kamat et al., 2016; Walker & Brown, 2018). Critically, the severity of apathy has strong implications for everyday functioning (e.g., Kamat et al., 2012; Kamat et al., 2013; Shapiro et al., 2013; Babicz et al., 2021). Given the profound functional consequences associated with apathy, conceptualizing the multidimensional neuropsychiatric syndrome, and associated neural mechanisms, following chronic HIV-1 viral protein exposure remains a critical need.

The neuropsychiatric syndrome of apathy is traditionally defined as a lack of motivation not attributable to other underlying impairments (Marin, 1991a); a definition which obfuscates objective measurement. The operational definition of apathy as a prominent decrease in self-generated voluntary or purposeful goal-directed behaviors (Levy & Dubois, 2006), however, has provided a method to quantify apathy. Goal-directed behaviors, which can range from single movements to complex behaviors, require the use of action to translate an internal state into the attainment of a goal. Achieving goal-directed behaviors requires several steps, including intention, initiation or execution, and evaluation (for review, Brown & Pluck, 2000); dysfunction at any of these steps may result in apathy. Accordingly, apathetic syndromes have been divided into three subtypes, including emotional-affective, cognitive, and auto-activation (Stuss et al., 2000; Levy & Dubois, 2006). Auto-activation apathy, specifically, is considered the most severe form of apathy and is characterized by a dramatic decrease in the self-initiation of spontaneous, voluntary behaviors (Levy & Dubois, 2006).

Voluntary wheel running, an experience that has naturally rewarding properties for rats (e.g., Kagan & Berkun, 1954; Collier & Hirsch, 1971; Greenwood et al., 2011; Basso & Morrell, 2015), affords one experimental paradigm for the evaluation of auto-activation apathy in preclinical biological systems. Under both natural (Meijer & Robbers, 2014) and laboratory (e.g., Stewart, 1898; Shirley, 1929; Basso & Morell, 2015) conditions, rodents voluntarily and spontaneously engage with a running wheel. At the macrostructural level (i.e., total running distance in one session), rodent wheel running is characterized by an acquisition and maintenance phase; phases which rely on goal-directed and habitual behavior, respectively (for review, Greenwood & Fleshner, 2019). Spontaneous activity (Richter, 1922), including voluntary wheel running (Eikelboom & Mills, 1988; Basso & Morell, 2017), however, occurs in distinct episodes that are organized into bouts of activity or disengagement. Parceling running behavior into fine-grained (e.g., one-minute) blocks using a microstructural analysis delineates the number and length of activity or disengagement bouts (Eikelboom & Mills, 1988). The relative importance of a microstructural analysis cannot be understated, as these analyses have been instrumental in elucidating motivational alterations resulting from neurotoxin exposure (Hoffman & Newland, 2016).

The neural circuits underlying goal-directed behaviors originate at the level of the brainstem (e.g., hypothalamus, periaqueductal gray) and incorporate brain regions (e.g., nucleus accumbens (NAc), prefrontal cortex (PFC)) associated with higher-order processing across phylogenetic and ontogenetic development (for review, Epstein & Silbersweig, 2015). The NAc, in particular, serves as the neural interface between the limbic and motor systems (Mogenson et al., 1980); an interface which results, at least in part, from the reception of a vast number of neurotransmitter inputs. Specifically, excitatory glutamatergic inputs from multiple cortical and limbic brain regions (e.g., amygdala, hippocampus, PFC, and thalamus; Phillipson & Griffiths, 1985) and dopaminergic projections from the ventral tegmental area (VTA; Fallon & Moore, 1978; Phillipson & Griffiths, 1985) converge on medium spiny neurons (MSNs) of the NAc. Experimental manipulations of either dopaminergic afferents to the NAc (Brown et al., 2012) or MSN function (Natsubori et al., 2017; Tsutsui-Kimura et al., 2017a; Tsutsui-Kimura et al., 2017b) strongly support the fundamental role of the NAc in goal-directed behaviors. Critically, chronic HIV-1 viral protein exposure induces decreased dopamine levels in the NAc (e.g., Jenuwein et al., 2004; Javadi-Paydar et al., 2017; Denton et al., 2019; for review, McLaurin et al., 2021a) and alters MSNs (Roscoe et al., 2014; Schier et al., 2017; McLaurin et al., 2018a; McLaurin et al., 2021b) supporting a potential neural mechanism underlying HIV-1 associated apathy.

Thus, the goals of the present study were threefold. First, to evaluate initiation, a key step for goal-directed behaviors and characteristic of auto-activation apathy, using voluntary wheel running behavior. Multiple variables, including time of day and wheel access, were systematically manipulated to establish the generality of deficits in the self-initiation of spontaneous behaviors. Second, to assess synaptic dysfunction in MSNs of the NAc following a history of voluntary wheel running. Third, to statistically establish synaptic dysfunction in the NAc as a neural mechanism of HIV-1 associated apathy. Establishing a fundamental neural mechanism underlying the self-initiation of spontaneous behavior affords a key target for the development of novel therapeutics and cure strategies for HIV-1 associated apathy.

## METHODS

### Experimental Design

A schematic of the experimental design, including assessments of goal-directed behaviors and synaptic dysfunction, is presented in **Figure 1A**.

**Figure 1.**
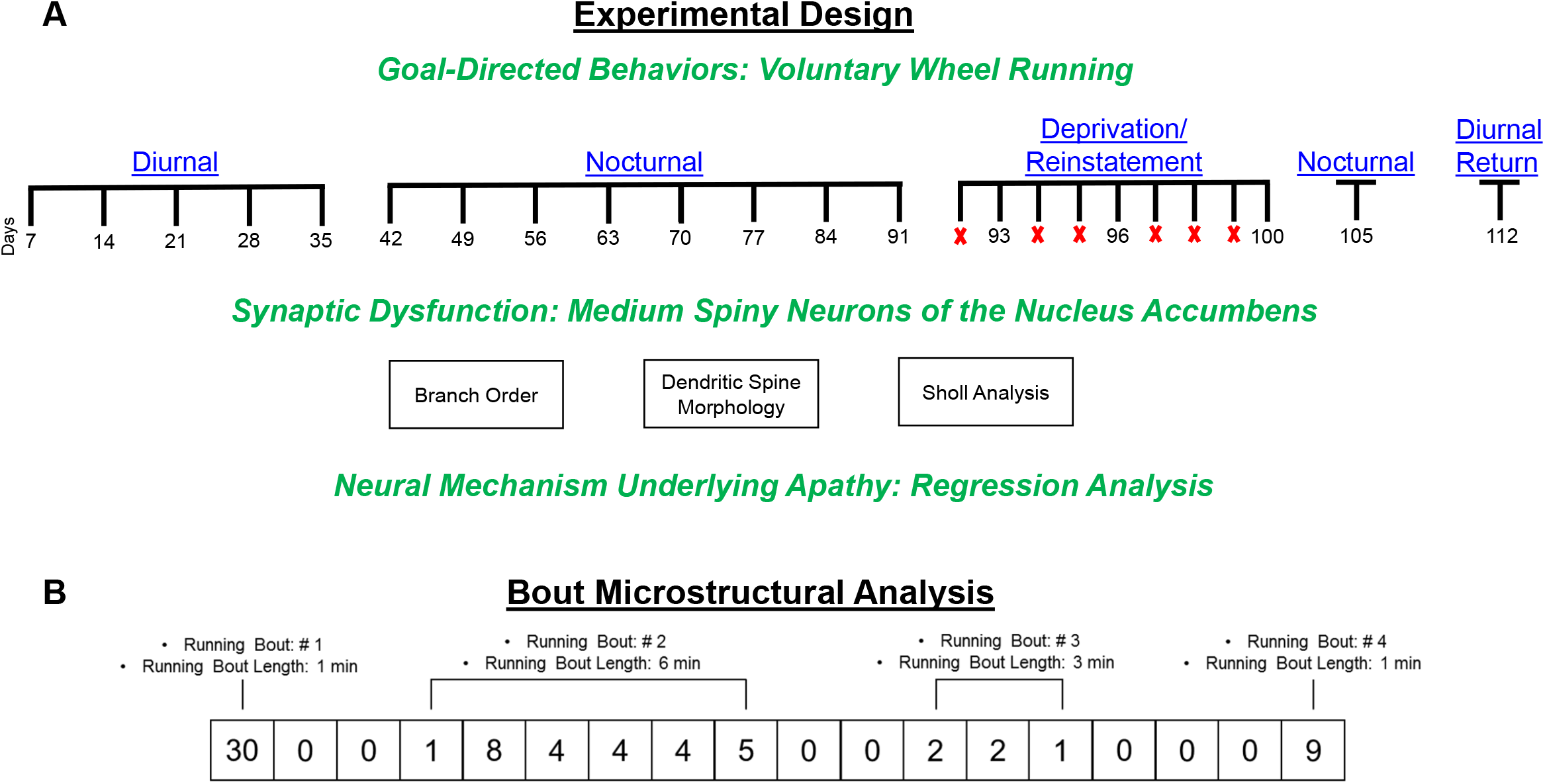
(**A**) Experimental Design Schematic (**B**) Illustration of the definitions (i.e., number of running bouts, running bout length) utilized for the bout microstructural analysis.

### Animals

Goal-directed behaviors, indexed using voluntary wheel running, were evaluated in ovariectomized (OVX) female Fischer (F344/N; Harlan Laboratories Inc., Indianapolis, IN) HIV-1 Tg (*n*=21) and control (*n*=26) animals. Prior to arrival at the animal facility, animals were OVX at Harlan Laboratories. Given that estrous cycle profoundly impacts voluntary wheel running behavior (e.g., Wang, 1923; Steiner et al., 1981; Basso & Morrell, 2017), female animals were OVX and fed a minimal phytoestrogen diet (≤20 ppm; Teklad 2020X Global Extruded Rodent Diet (Soy Protein-Free)). HIV-1 Tg and control animals were delivered to the animal vivarium between postnatal day (PD) 45 and PD 60 and pair- or group-housed throughout the duration of the experiment. Rodent food and water were available *ad libitum*.

Animals were maintained in AAALAC-accredited facilities using the guidelines established by the National Institutes of Health in the Guide for the Care and Use of Laboratory Animals. Conditions of the animal colony were targeted at 21° ± 2° C, 50% ± 10% relative humidity and had a 12-h light: 12-h dark cycle with lights on at 0700 h (EST). The project protocol was approved by the Institutional Animal Care and Use Committee at the University of South Carolina (Federal Assurance: #D16-00028).

### Experimental Controls

#### Somatic Growth: Body Weight

Body weight, an index of somatic growth, was measured five days per week for the duration of the study.

#### Integrity of Gross Motoric System Function: Locomotor Activity

##### Apparatus

Perspex inserts were utilized to convert square (40 x 40 cm) activity monitors (Hamilton Kinder, San Diego, CA) into round (~40 cm diameter) compartments. Free motor movement was detected using infrared photocells (32 emitter/detector pairs) which were spaced approximately 2.5 cm apart. The manufacturer tuned the infrared photocells to maintain their sensitivity with the additional perspex insert.

##### Procedure

Locomotor activity, an index of gross motoric system function, was evaluated weekly for seven consecutive weeks. Assessments were conducted using a 60-min test session from 1200 h to 1600 h in an isolated room. Dim light conditions (<10 lx) and background white noise (70 dB(A)) were present throughout the duration of testing. Gross motoric system function was evaluated using the total number of photocell interruptions within the 60-min test session.

### Goal-Directed Behaviors: Voluntary Wheel Running

#### Apparatus

A 34 cm diameter running wheel was attached to the lid of a polypropylene rectangular chamber (45.5cm long, 24cm wide and 20.5cm deep). Wheel revolutions were recorded by magnetic sensors, which were integral to the lid and wheel, by Vital View 4.0 (Mini Mitter Inc., Bend, OR). The number of wheel revolutions during each minute of testing was collected for analysis.

#### Procedure

##### Diurnal Voluntary Wheel Running

Voluntary wheel running in HIV-1 Tg and control animals was evaluated during the diurnal phase (i.e., 1200 h to 1700 h) of their light-dark cycle. Animals had daily access to the running wheel apparatus, which was located in an isolated room with low-light conditions (<10 lx), for approximately 66-min. Diurnal running was continued until the overall genotype mean for the total number of meters run was stabilized for three consecutive weeks. In total, the diurnal voluntary wheel running phase was 35 days (5 weeks).

##### Nocturnal Voluntary Wheel Running

After reaching stability during the diurnal phase of their light-dark cycle, HIV-1 Tg and control animals were assessed during the nocturnal phase (i.e., 19:00 h to 2400 h) using the procedures and criteria described above. The nocturnal voluntary wheel running phase was 56 days (8 weeks).

##### Deprivation/Reinstatement

Following three weeks of stabilized nocturnal voluntary wheel running behavior, running wheels were locked to prevent rotation in a dose-response manner (i.e., 1, 2, or 3 days). During nocturnal deprivation, HIV-1 Tg and control animals were placed in the apparatus with a locked running wheel for approximately 66-min. Reinstatement responses were recorded by providing access to the running wheel for one day after each wheel-locking dose. The number of days the wheels were locked was presented in an ascending manner.

##### Nocturnal Voluntary Wheel Running

After the final reinstatement response recording, nocturnal voluntary wheel running assessments were conducted for an additional five days.

##### Diurnal Return

The overarching goal of the diurnal return was to determine whether differences in the average number of meters run per day during the diurnal and nocturnal phases were due to learning or entrainment to the circadian cycle. HIV-1 Tg and control animals were returned to a diurnal voluntary wheel running schedule for 1 week (7 days).

#### Bout Microstructural Analysis

A bout microstructural analysis was conducted by parceling running behavior into 1-min blocks of activity (i.e., running bout) or disengagement using methodology similar to previous reports (Eikelboom & Mills, 1988; **Figure 1B**). A running bout was defined as the performance of at least one wheel revolution during consecutive one-min bins. Two key components of running bouts were extrapolated, including the total number of running bouts in a session and the average running bout length.

### Synaptic Dysfunction: Medium Spiny Neurons of the Nucleus Accumbens

#### Tissue Preparation

After the completion of their final voluntary wheel running assessment, HIV-1 Tg and control animals were deeply anesthetized with sevoflurane (Abbot Laboratories, North Chicago, IL) and transcardially perfused using the methodology reported by Roscoe et al. (2014) with two minor modifications. Specifically, post-fixation occurred in 4% chilled paraformaldehyde for 10 minutes and 1000 μm thick coronal sections were cut using the rat brain matrix (ASI Instruments, Warren, MI).

#### Preparation of Tefzel Tubing and DiI Cartridges

A polyvinylpyrrolidone (PVP; Sigma-Aldrich, St. Louis, MO) solution was prepared by dissolving 100 mg of PVP in ddH_2_O. The PVP solution was drawn into Tefzel tubing (70 cm; IDEZ Health Sciences, Oak Harbor, WA), allowed to coat for at least 10 minutes, and was removed.

DiOlisitic cartridges were prepared by separately dissolving 175 mg of tungsten beads (Bio-Rad, Hercules, CA) and 7 mg crystallized DiI in 99.5% methylene chloride. After applying the tungsten bead solution to a glass slide, the DiI solution was titrated on top; the two solutions were mixed slowly and allowed to completely dry. The dye/bead mixture was scraped into two 1.5 mL centrifuge tubes with 1.5 mL ddH_2_O and sonicated until homogenous. The homogenized mixture from the two 1.5 mL centrifuge tubes was combined in a 15 mL conical tube and sonicated further before being drawn into the PVP coated Tefzel tubing. Once filled with the homogenized tungsten beads-DiI solution, the Tefzel tubing was loaded into the tubing prep station (Bio-Rad, Hercules CA) and rotated for 5 minutes. The remaining water was slowly drawn from the tubing. Nitrogen gas was turned on at a flow rate of 0.4 L/min and the tubing was rotated for an additional 30 minutes. Tubing was cut into 13 mm lengths and stored in the dark.

#### DiOlistic Labeling and Confocal Imaging

MSNs from the NAc, located approximately 3.24 mm to 0.48 mm anterior to Bregma (Paxinos & Watson, 2014) were ballistically labeled and mounted using the procedures reported by Roscoe et al. (2014). For each animal, multiple (i.e., 3-5) Z-stack images of the DiOlistically labeled neurons were taken using a Nikon TE-2000E and Nikon’s EZ-C1 computing software (v. 3.81b). Sholl analysis, dendritic branching complexity, and dendritic spine morphology analyses were conducted on one neuron from each animal; neurons were blinded to prevent experimenter bias. Neurons included in the analysis were selected based on established selection criteria (Li et al., 2020), yielding Control, *n*=19 and HIV-1 Tg, *n*=12. Selected neurons were analyzed using the AutoNeuron and AutoSpine extension modules within Neurolucida360 (MicroBrightfield, Williston, VT).

#### Neuronal and Dendritic Spine Analyses

Morphological parameters of MSNs in the NAc were examined using two complementary measures, namely Sholl analysis (Sholl, 1953) and dendritic branch ordering. For the Sholl analysis, concentric circles were placed at 10 μm radius intervals and the number of intersections at each successive radii were measured. For dendritic branch ordering, the number of segments at branch orders 1 through 10 were evaluated; branch orders were assigned using a centrifugal branch ordering method.

Dendritic spine morphology was evaluated using continuous measurements of three parameters, including backbone length (μm), head diameter (μm) and neck diameter (μm). Dendritic spines were included in the analysis based on previously reported results (i.e., backbone length, 0.2 to 4.0 μm (Fernández-Cabrera et al., 2018); head diameter, 0.0 to 1.2 μm (Shen et al., 2008); volume, 0.05 to 0.85 μm^3^ (for review, Harris & Kater, 1994).

### Statistical Analysis

Analysis of variance (ANOVA) and regression techniques were utilized for the statistical analysis of all data (SAS/STAT Software 9.4, SAS Institute Inc., Cary, NC; SPSS Statistics 27, IBM Corp., Somer, NY; GraphPad Prism 5 Software Inc., La Jolla, CA). Potential violations of sphericity were precluded by the use of either orthogonal decompositions or the conservative Greenhouse-Geisser *df* correction factor (*p*_GG_; Greenhouse & Geisser, 1959). Statistical significance was evaluated at a *p* value of 0.05.

For ANOVA analyses, genotype (HIV-1 Tg vs. control) served as the between-subjects factor, while time (e.g., age, weeks, day), radius, branch order, or bin served as the within-subjects factor, as appropriate. For the deprivation/reinstatement phase, the dependent variable of interest after either one, two, or three days of wheel locking were compared to the average total number of meters run during week 13 (i.e., the final week of nocturnal voluntary wheel running cooresponding to Day 91).

Experimental controls, including both body weight and locomotor activity, were analyzed using a mixed-model ANOVA (SPSS Statistics 27) and regression techniques (GraphPad Prism 5 Software, Inc.).

At the macrostructural level, the dependent variable of interest was the total number of meters run. The total number of meters run during an individual daily session were averaged across seven days for the statistical analysis. A mixed-model ANOVA (SPSS Statistics 27) and complementary regression techniques (GraphPad Prism 5 Software, Inc.) were conducted to analyze voluntary wheel running behavior at the macrostructural level.

At the microstructural level, two dependent variables, including running bout number and running bout length, were investigated. The number of running bouts and running bout length during an individual daily session were averaged across seven sessions for the statistical analysis. Running bout number during the diurnal, nocturnal, and deprivation/reinstatement phases were analyzed using a generalized linear mixed effects model with a Poisson regression and unstructured covariance structure (PROC GLIMMIX; SAS/STAT Software 9.4). For the diurnal voluntary wheel running phase, the number of running bouts were squared to transform the data. Running bout lengths were analyzed using a mixed-model ANOVA (SPSS Statistics 27). Additionally, complementary regression analyses were conducted to evaluate the functional form of the cumulative frequency of running bout number and running bout length.

Genotypic alterations in neuronal and dendritic spine morphology were examined to evaluate synaptic dysfunction. The number of intersections at successive radii, a dependent variable derived from the Sholl analysis, was analyzed using a mixed-design ANOVA with a variance components covariance structure, a random intercept, and a random slope (i.e., Radius; Wilson et al., 2017; PROC MIXED; SAS/STAT Software 9.4). The number of dendritic branches at each successive branch order, the number of dendritic spines between radii, and morphological characteristics of dendritic spines (i.e., Backbone Length, Head Diameter, Neck Diameter) were examined using a generalized linear mixed effects model with a Poisson regression and unstructured covariance structure (PROC GLIMMIX; SAS/STAT Software 9.4). Notably, only dendritic spines with a head diameter greater than 0.01 μm were included in the figure and analysis of dendritic spine head diameter.

Multiple linear regression analyses (SPSS Statistics 27, IBM Corp., Somer, NY; GraphPad Software, Inc., La Jolla, CA) were conducted to evaluate synaptic dysfunction as a potential neural mechanism underlying self-initiation of spontaneous behaviors. The three dependent variables of interest were the average number of running bouts, an index of the self-initiation of spontaneous behaviors, during the diurnal, nocturnal, and deprivation/reinstatement phases. Independent variables included measures of synaptic dysfunction that exhibited a statistically significant genotype effect (i.e., Number of Dendritic Spines Between Radii, Relative Frequency of Dendritic Branches at Each Branch Order, Number of Dendritic Spines Within Each Bin for Backbone Length, Head Diameter, and Neck Diameter).

### Data Availability Statement

All relevant data are within the paper.

## RESULTS

### Experimental Controls

#### Somatic Growth: Body Weight

Independent of genotype, animals exhibited a significant increase in body weight throughout development; a developmental trajectory that was well-described by a one-phase association (**Supplementary Figure 1A**; *R*^2^s ≥ 0.98). Although HIV-1 Tg animals weighed significantly less than control animals throughout the experiment (Main Effect: Genotype, *F*(1,45)=29.9, *p*≤ 0.001, η_p_^2^=0.400), there was no statistically significant difference in the rate of somatic growth (Genotype x Age Interaction, *p*>0.05).

#### Integrity of Gross-Motoric System Function: Locomotor Activity

HIV-1 Tg and control animals exhibited intact gross-motoric system function, indexed using a 60-min locomotor activity test session for seven consecutive weeks. Independent of genotype, the total number of photocell interruptions was well-described by a sigmoidal dose-response function (**Supplementary Figure 1B**; *R*^2^s ≥ 0.95). Despite the downward mean shift in the total number of photocell interruptions observed in HIV-1 Tg animals (Main Effect: Genotype, [*F*(1,45)=26.7, *p*≤0.001, η_p_^2^=0.372]), the number of photocell interruptions were significantly greater than zero across all testing sessions (i.e., H_0_: Bottom of the Sigmoidal Dose Response Curve = 0; Control: [*F*(1,179)=19.8, *p*≤0.001], HIV-1 Tg: [*F*(1,144)=20.0, *p*≤0.001]). Consistent with previous reports (McLaurin et al., 2018b), therefore, neither HIV-1 Tg nor control animals exhibited any gross-motoric system impairments.

### Goal-Directed Behaviors: Voluntary Wheel Running

After establishing the integrity of gross-motoric system function, goal-directed behaviors, an index of apathy, were evaluated using voluntary wheel running. HIV-1 Tg and control animals were given daily access to a running wheel for approximately 66-min during both the diurnal and nocturnal phases.

#### Diurnal Voluntary Wheel Running

During the diurnal phase, presence of the HIV-1 transgene failed to alter (*p*>0.05) running behavior at the macrostructural level (i.e., total running distance in one session averaged across seven sessions; **Figure 2A**). The average number of meters run per day was well-described by a global one-phase decay (*R*^2^ = 0.82), whereby both HIV-1 Tg and control animals exhibited an initial decrease, and subsequent plateau, in the average daily running distance (Main Effect: Week, [*F*(4,180)=46.1, *p*_GG_≤0.001, η_p_^2^=0.506] with a prominent linear component [*F*(1,45)=78.6, *p*≤0.001, η_p_^2^=0.636]).

**Figure 2.**
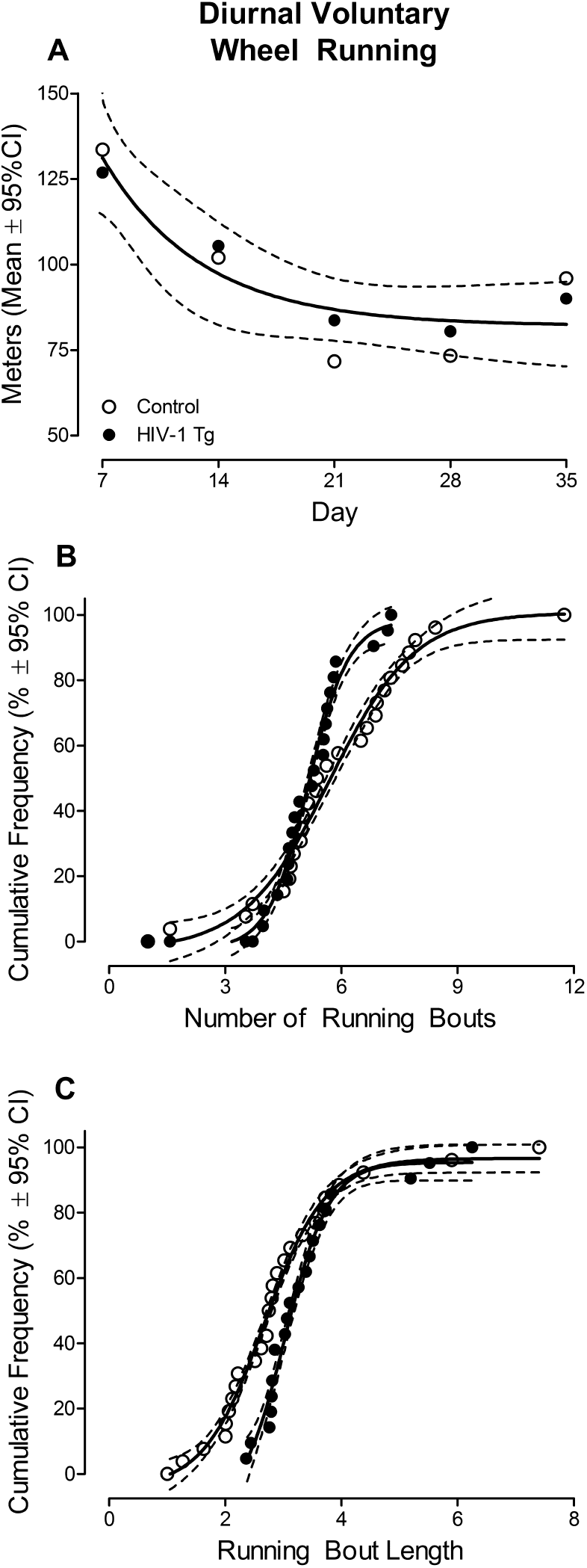
Motivation for diurnal voluntary wheel running at the macrostructural (i.e., total number of meters run in one session averaged across seven days; **A**) and microstructural (i.e., number of running bouts, running bout length; **B-C**) level is illustrated as a function of genotype (HIV-1 Tg vs. Control; ±95% Confidence Intervals). At the macrostructural level, presence of the HIV-1 transgene failed to alter running distance (**A**). At the microstructural level, HIV-1 Tg animals exhibited prominent motivational alterations, evidenced by a profound decrease in the number of running bouts, relative to control animals (**B**); observations which support deficits in the self-initiation of spontaneous behaviors. However, once wheel running behavior was initiated, HIV-1 Tg animals displayed a longer running bout length relative to control animals (**C**).

Prominent motivational alterations, however, were revealed by a bout microstructural analysis, which characterized the pattern of voluntary wheel running behavior. A population shift towards a decreased number of running bouts was observed in HIV-1 Tg animals relative to controls (**Figure 2B**); the magnitude, but not pattern, of which was dependent upon week (Genotype x Week Interaction, [*F*(4,180)=8.0, *p*≤0.001]). Complementary regression analyses, therefore, were conducted on the average number of running bouts during the diurnal phase (i.e., collapsed across week). Independent of genotype, the average number of running bouts during the diurnal phase were well-described by a sigmoidal dose-response curve with a variable slope (Control: *R*^2^=0.97; HIV-1 Tg: *R*^2^=0.98); albeit significant differences in the parameters of the function were observed [*F*(4,44)=26.7, *p*≤0.001].

With regards to running bout length, HIV-1 Tg animals exhibited a population shift towards longer running bouts relative to control animals (**Figure 2C**); the magnitude, but not pattern, of which was, again, dependent upon week (Genotype x Week Interaction with a prominent linear component [*F*(1,45)=5.7, *p*≤0.02, η_p_^2^=0.113]). The average running bout length during the diurnal phase (i.e., collapsed across week) was utilized to conduct complementary regression analyses. Specifically, a sigmoidal dose-response curve with a variable slope afforded the best-fit for the cumulative frequency of the average running bout length during the diurnal phase (Control: *R*^2^=0.98; HIV-1 Tg: *R*^2^=0.98). However, significant genotypic differences in the parameters of the function were observed [*F*(4,38)=57.3, *p*≤0.001].

Taken together, during the diurnal phase, motivational alterations in the HIV-1 Tg rat were evidenced at the microstructural, but not macrostructual, level of analysis. HIV-1 Tg animals exhibited a profound decrease in the number of running bouts relative to control animals supporting a deficit in the self-initiation of spontaneous behaviors. Notably, however, once wheel running behavior was initiated, HIV-1 Tg animals displayed a longer running bout length.

#### Nocturnal Voluntary Wheel Running

During the nocturnal phase, prominent motivational alterations were evidenced at the macrostructural level by a genotype-dependent development of voluntary wheel running behavior across time (**Figure 3A**; Genotype x Week Interaction, [*F*(7,308)=3.0, *p*_GG_≤0.04, η_p_^2^=0.064] with a prominent cubic component [*F*(1,44)=4.7, *p*≤0.04, η_p_^2^=0.096]). In control animals, the average number of meters run per session was well-described by a sigmoidal dose response curve (*R*^2^=0.95), whereas a segmental linear regression (*R*^2^=0.85) afforded the best-fit for HIV-1 Tg animals. Specifically, during the acquisition phase, HIV-1 Tg animals increased their average daily running distance, faster than control animals. After voluntary wheel running behavior stabilized (i.e., the maintenance phase), however, control animals exhibited a significantly greater average number of meters run per session relative to HIV-1 Tg animals.

**Figure 3.**
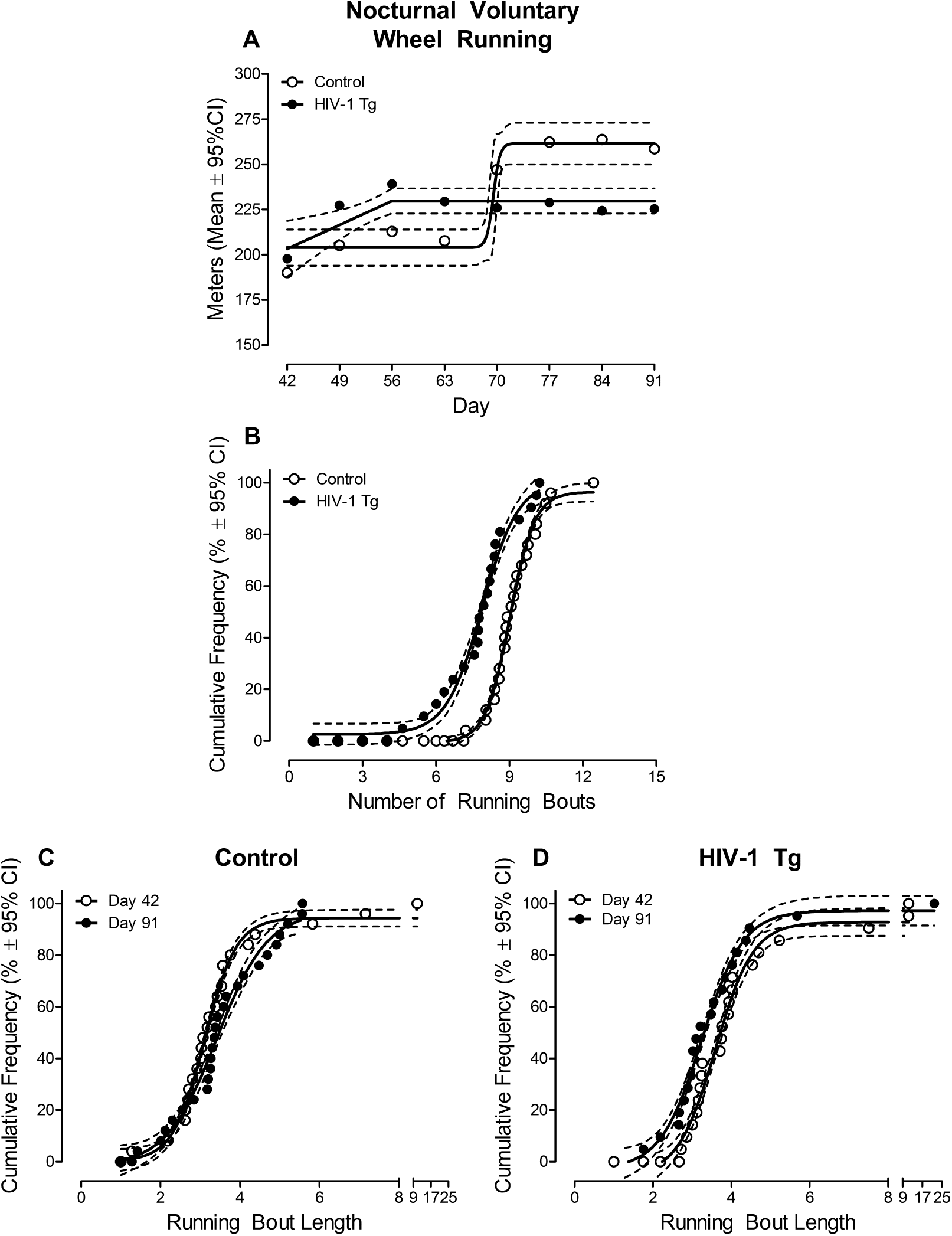
Observations during the nocturnal voluntary wheel running phase are illustrated as a function of genotype (HIV-1 Tg vs. Control; ±95% Confidence Intervals). At the macrostructural level (i.e., total number of meters run in one session averaged across seven days) presence of the HIV-1 transgene altered the development of voluntary wheel running behavior across time (**A**). With regards to the number of running bouts, HIV-1 Tg animals exhibited a prominent population shift towards fewer running bouts relative to control animals supporting a deficit in the self-initiation of spontaneous behaviors (**B**). Furthermore, prominent alterations in the development of running bout length were observed in HIV-1 Tg animals relative to control animals (**C-D**).

Motivational alterations in the HIV-1 Tg rat were also observed at the microstructural level, characterized by prominent genotypic differences in the pattern of nocturnal voluntary wheel running behavior. HIV-1 Tg animals exhibited a population shift towards fewer running bouts, independent of week, relative to control animals (**Figure 3B**; Main Effect: Genotype, *F*(1,44)=14.8, *p*≤0.001). Complementary regression analyses, collapsed across week, further confirmed the observations, whereby a sigmoidal dose-response curve with a variable slope afforded a well-described function for the number of running bouts in both HIV-1 Tg (*R*^2^=0.98) and control (*R*^2^=0.99) animals; albeit significant differences in the parameters of the function were observed [*F*(4,50)=227.3, *p*≤0.001].

Furthermore, presence of the HIV-1 transgene altered the development of running bout length across time (**Figure 3C-D**; Genotype x Week Interaction with a prominent linear component, [*F*(1,44)=4.5, *p*≤0.04, η_p_^2^=0.092]). Complementary regression analyses were conducted at Day 42 and 91, representing the first and last weeks of nocturnal voluntary wheel running, respectively. In control animals, a statistically significant population shift towards longer running bouts was observed across time (Best Fit Function: Sigmoidal Dose-Response with a Variable Slope, *R*^2^=0.99 (Day 42) and *R*^2^=0.97 (Day 91); Differences in the Parameters of the Function: [*F*(4,43)=20.1, *p*≤0.001]). HIV-1 Tg animals, in sharp contrast, exhibited a prominent population shift towards shorter running bouts across time; running bout length at both Day 42 (*R*^2^=0.97) and Day 91 (*R*^2^=0.98) were well-described by a sigmoidal dose-response curve with a variable slope (Differences in the Parameters of the Function: [*F*(4,39)=30.3, *p*≤0.001]).

Collectively, during the nocturnal phase, motivational alterations at the macrostructural level were characterized by alterations in both the acquisition and maintenance phase of voluntary wheel running in the HIV-1 Tg rat. Examination of the bout microstructure revealed a profound decrease in the number of running bouts and prominent alterations in the development of running bout length in HIV-1 Tg animals relative to control animals. Deficits in the self-initiation of spontaneous behaviors, therefore, generalize across both the diurnal and nocturnal phases.

#### Deprivation/Reinstatement

After exhibiting a stabilized running distance for at least three weeks during the maintenance phase of nocturnal voluntary wheel running, HIV-1 Tg and control animals were deprived of their running wheels for one, two, and three days; a dose response paradigm that was conducted in an ascending manner. Reinstatement responses were recorded by providing access to the running wheel for one day after each wheel-locking dose.

At the macrostructual level, both HIV-1 Tg and control animals exhibited a profound burst of rebound running relative to stabilized total running distances at Day 91 (i.e., the final week of the nocturnal voluntary wheel running phase; **Figure 4A**; Main Effect: Deprivation, [*F*(3,132)=3.7, *p*_GG_≤0.02, η_p_^2^=0.078] with a prominent linear component [*F*(1,44)=6.1, *p*≤0.02, η_p_^2^=0.078]). However, neither presence of the HIV-1 transgene nor deprivation dose altered the rebound running response at the macrostructural level (*p*>0.05).

**Figure 4.**
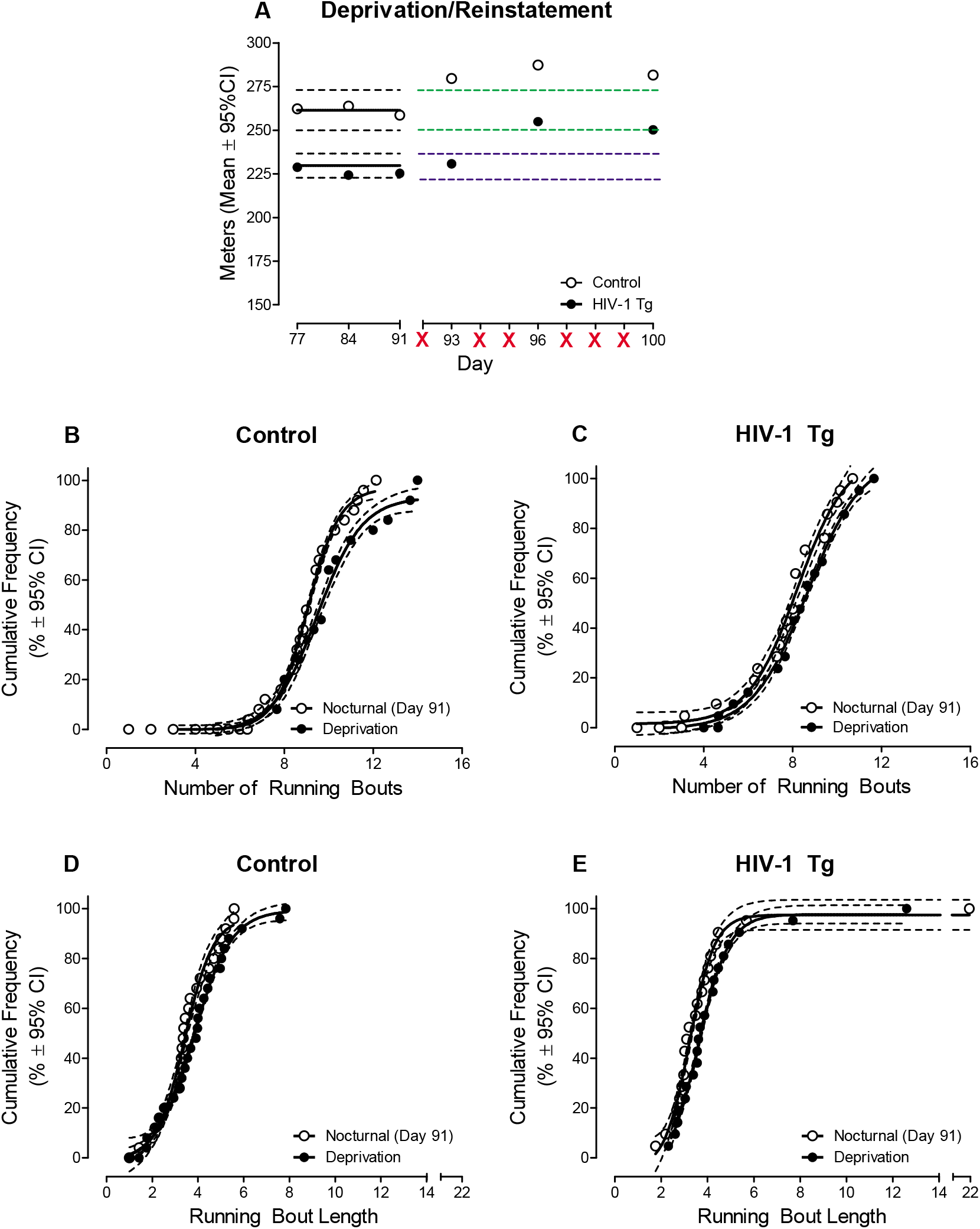
After exhibiting a stabilized running distance for at least three weeks during the maintenance phase of nocturnal voluntary wheel running, HIV-1 Tg and control animals were deprived of their running wheels for one, two, and three days. Reinstatement responses, illustrated as a function of genotype (HIV-1 Tg vs. Control; ±95% Confidence Intervals), were examined by providing access to the running wheel for one day after each wheel-locking dose. No prominent genotypic alterations in the reinstatement response were observed at either the macrostructural (**A**) or microstructural (**B-E**) level. In panel **A**, the green (Control) and blue (HIV-1 Tg) dashed lines in reflect an extension of the 95% confidence interval from the best fit function during the nocturnal voluntary wheel running session.

Similar observations were observed at the microstructural level. First, with regards to the number of running bouts, HIV-1 Tg animals exhibited a decreased bout number, independent of deprivation, relative to control animals (Main Effect: Genotype, [*F*(1,44)=9.1, *p*≤0.001]). Second, neither presence of the HIV-1 transgene nor deprivation significantly altered running bout length during the rebound phase (*p*>0.05). Collectively, examination of reinstatement responses following maintenance of nocturnal voluntary wheel running revealed no prominent genotypic alterations.

### Synaptic Dysfunction: Medium Spiny Neurons of the Nucleus Accumbens

#### Neuronal Morphology

HIV-1 Tg animals exhibited selective alterations in the morphological characteristics of MSNs of the NAc; characteristics which were evaluated using two complementary measures, namely Sholl analysis and centrifugal branch ordering. Presence of the HIV-1 transgene failed to significantly alter dendritic arbor complexity (**Figure 5A**; Genotype x Radius Interaction: *p*>0.05), indexed using Sholl analysis, evidenced by a global best fit function (i.e., Sum of Two Gaussians, *R*^2^=0.97). With regards to centrifugal branch ordering (**Figure 5B**), however, HIV-1 Tg animals displayed a prominent rightward shift with an increased relative frequency of higher order dendritic branches relative to control animals (Genotype x Branch Order Interaction: [*F*(9, 261)=6.4, *p*≤0.001]). The number of dendritic branches at each successive branch order were well-described by a Gaussian fit for both HIV-1 Tg (*R*^2^=0.94) and control (*R*^2^=0.98) animals; albeit significant differences in the parameters of the function were observed [*F*(3,14)=15.3, *p*≤0.001]. HIV-1 Tg animals, therefore, exhibited enhanced dendritic branching complexity, evidenced via centrifugal branch ordering, relative to control animals.

**Figure 5.**
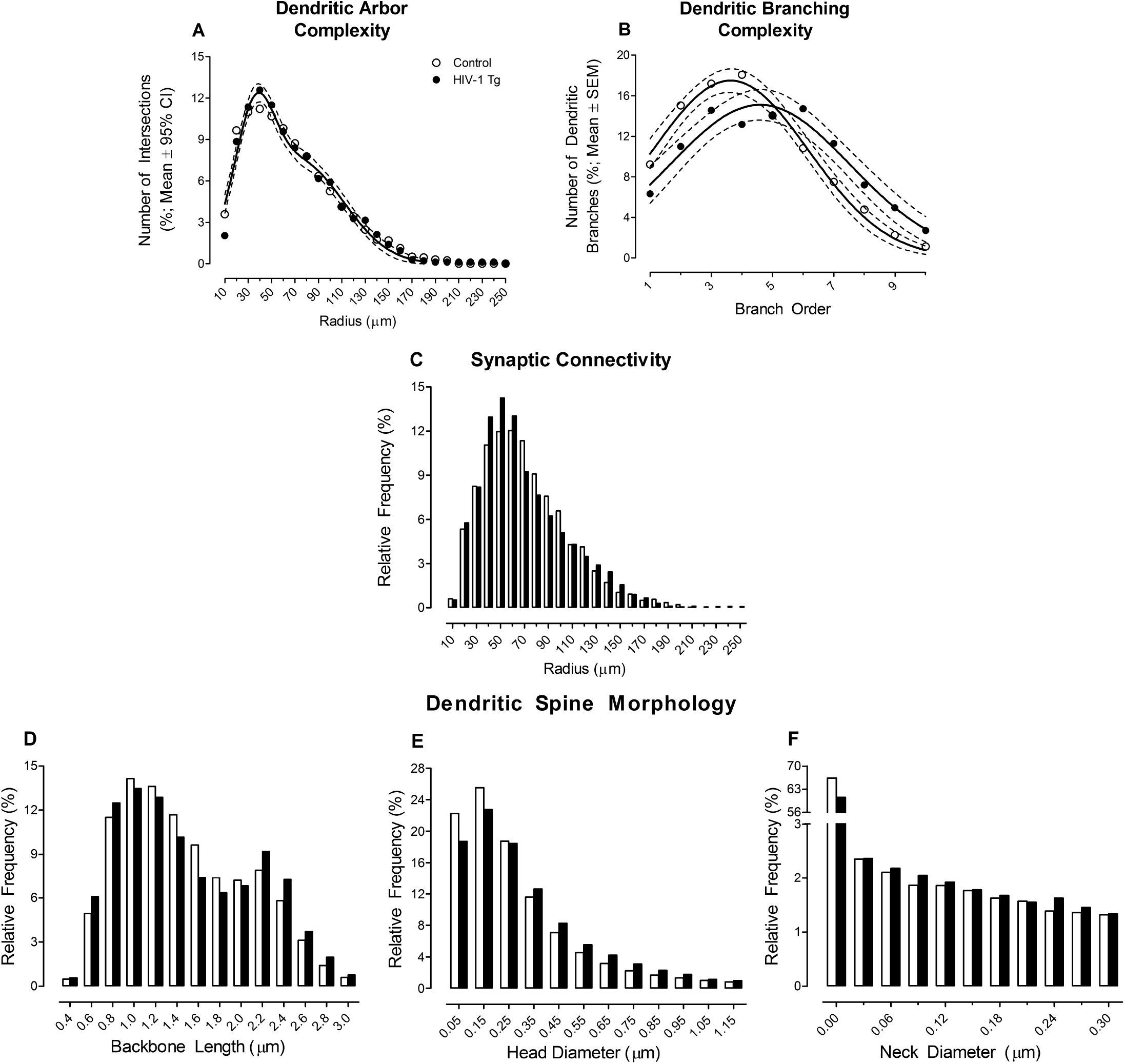
Following the completion of voluntary wheel running, the influence of the HIV-1 transgene on neuronal morphology (**A-B**; ±95% Confidence Intervals), synaptic connectivity (**C**) and dendritic spine morphology (**D-F**) in medium spiny neurons in the nucleus accumbens was examined. Data are illustrated as a function of genotype (HIV-1 Tg vs. Control). HIV-1 Tg animals exhibited selective alterations in neuronal morphology, characterized by enhanced dendritic branching complexity (**B**); albeit presence of the HIV-1 transgene failed to significantly alter dendritic arbor complexity (**A**). Deficits in synaptic connectivity were evidenced by an increased relative frequency of dendritic spines on branches more proximal to the soma in HIV-1 Tg animals relative to control animals (**C**). The morphological parameters of dendritic spines in HIV-1 Tg animals were characterized by a bimodal distribution for dendritic spine backbone length (**D**) and a population shift towards increased dendritic spine head (**E**) and neck diameter (**F**) relative to control animals; observations which support a population shift towards a ‘stubby’ dendritic spine phenotype in HIV-1 Tg animals.

#### Synaptic Connectivity

The location of dendritic spines on MSNs of the NAc, an important factor given that proximal and distal dendrites receive different afferents (for review, Spiga et al., 2014), were analyzed using Sholl analysis. HIV-1 Tg animals exhibited an increased relative frequency of dendritic spines on branches more proximal to the soma relative to control animals (**Figure 5C**; Genotype x Radius Interaction, [*F*(21,696)=25.9, *p*≤0.001]) supporting an alteration in synaptic connectivity.

#### Dendritic Spine Morphology

Constitutive expression of the HIV-1 transgene shifted the morphological parameters of dendritic spines towards an immature phenotype (**Figure 5D-F**). Presence of the HIV-1 transgene altered the distribution of dendritic spine backbone length relative to control animals (**Figure 5D**; Genotype x Bin Interaction, [*F*(13,377)=18.2, *p*≤0.001]). Specifically, the distribution of dendritic spine backbone length in HIV-1 Tg animals was characterized by a bimodal distribution, whereby they exhibited increased relative frequencies of dendritic spines at the ends of the distribution relative to control animals. Additionally, HIV-1 Tg animals exhibited a population shift towards dendritic spines with increased head diameter (**Figure 5E**; Genotype x Bin Interaction, [*F*(11,319)=15.7, *p*≤0.001]) and increased neck diameter (**Figure 5F**; Genotype x Bin Interaction, [*F*(10,290)=3.1, *p*≤0.001]) relative to control animals. Taken together, morphological parameters of dendritic spines in HIV-1 Tg animals support a population shift towards a ‘stubby’ dendritic spine phenotype relative to control animals.

### Neural Mechanism Underlying Apathy: Regression Analyses

Synaptic dysfunction mechanistically underlies the self-initiation of spontaneous behaviors, indexed using voluntary wheel running, during the diurnal, nocturnal, and deprivation/reinstatement phases. During the diurnal phase (**Figure 6A**), regressing the relative frequency of dendritic branches at Branch Order 3, Branch Order 10, and Branch Order 8 on the number of running bouts revealed a regression coefficient (*r*) of 0.648. Changes in neuronal morphology, therefore, explained 42.0% of the variance in the number of running bouts during the diurnal phase. During the nocturnal phase (**Figure 6B**), seven variables reflecting measures of neuronal morphology (Relative Frequency of Dendritic Branches at Branch Order 7 and Branch Order 2), dendritic spine morphology (Number of Dendritic Spines in Head Diameter Bin 0.85 μm, Head Diameter Bin 1.05 μm, Backbone Length Bin 2.2 μm, and Head Diameter Bin 0.95 μm) and synaptic connectivity (Number of Dendritic Spines at Radius 30) explained 68.5% of the variance (*r*=0.827) in the number of running bouts. During the deprivation/reinstatement phase (**Figure 6C**), 54.5% of the variance (*r*=0.738) in the number of running bouts was explained by six variables reflecting measures of neuronal morphology (Relative Frequency of Dendritic Branches at Branch Order 1 and Branch Order 5), dendritic spine morphology (Number of Dendritic Spines in Head Diameter Bin 0.95 μm and Neck Diameter Bin 0.21 μm) and synaptic connectivity (Number of Dendritic Spines at Radius 110 and Radius 20). Regression analyses, therefore, support synaptic dysfunction as a fundamental neural mechanism underlying the self-initiation of spontaneous behavior, a key step for goal-directed behaviors.

**Figure 6.**
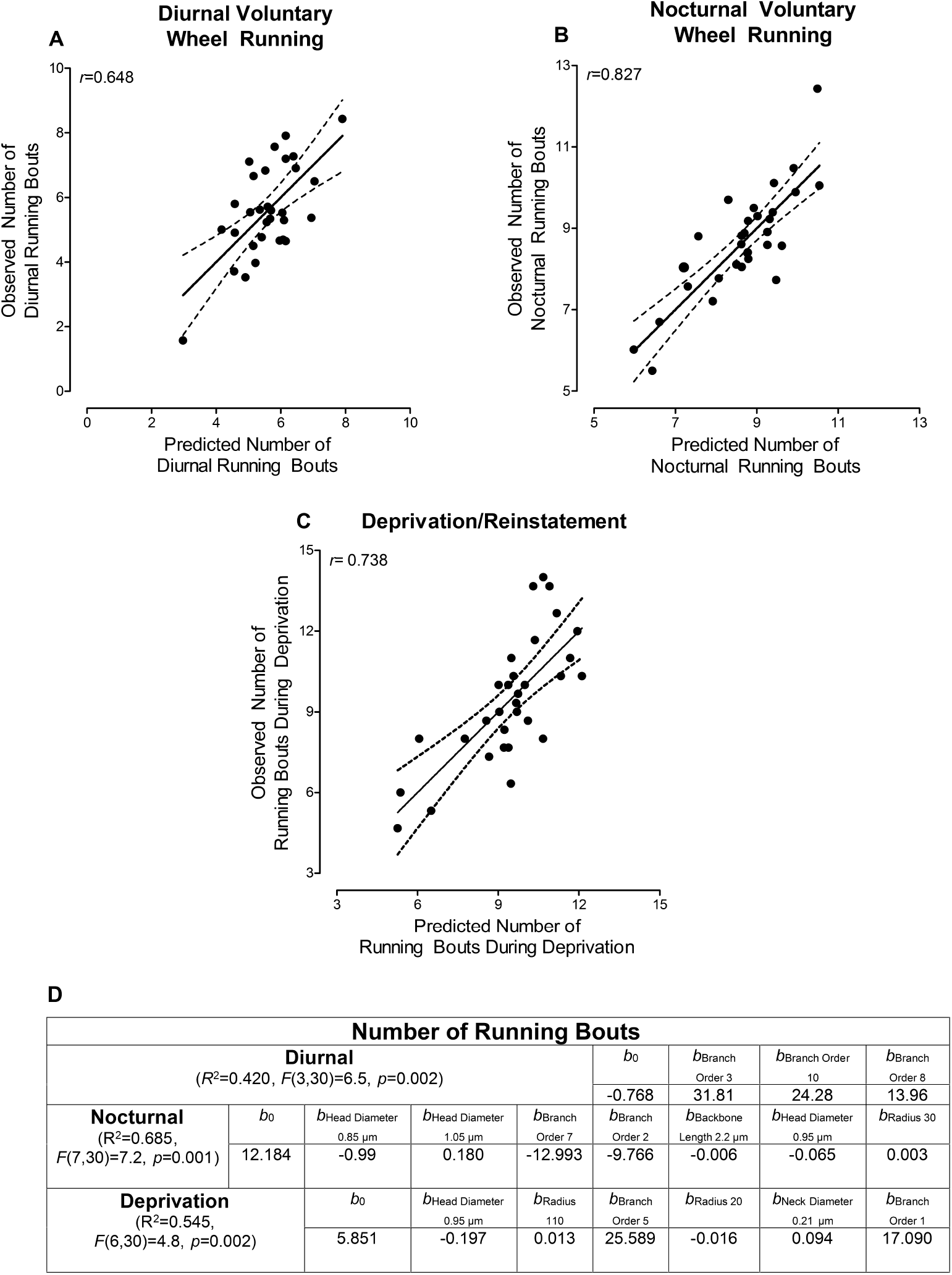
Synaptic dysfunction mechanistically underlies the self-initiation of spontaneous behaviors during the diurnal phase (**A**), nocturnal phase (**B**), and deprivation/reinstatement phase (**C**). During the diurnal phase, measures of neuronal morphology explained 42.0% of the variance in the number of running bouts (**A**). During the nocturnal phase, measures of synaptic dysfunction explained 68.5% of the variance in the number of running bouts (**B**). During the deprivation/reinstatement phase, 54.5% of the variance in the number of running bouts was explained by measures of synaptic dysfunction (**C**). Panel (**D**) reports the results and coefficients from the multiple regression analysis.

## DISCUSSION

Profound alterations in purposeful goal-directed behaviors, an index of apathetic behavior, were observed in HIV-1 Tg animals using voluntary wheel running. Apathetic behaviors in the HIV-1 Tg rat were characterized by deficits in the self-initiation of spontaneous behaviors, evidenced by a prominent decrease in the number of running bouts relative to control animals; alterations which persisted during both the diurnal and nocturnal cycle. Following the completion of voluntary wheel running, MSNs of the NAc were examined as a potential neural mechanism of apathetic behaviors in the HIV-1 Tg rat. Synaptic dysfunction in MSNs of the NAc of HIV-1 Tg animals was characterized by decreased synaptic connectivity and a population shift towards an immature dendritic spine phenotype relative to control animals. Multiple regression analyses established synaptic dysfunction as a key neural mechanism underlying the self-initiation of spontaneous behaviors. Taken together, synaptic dysfunction may afford a key target for the development of novel therapeutics and cure strategies for HIV-1 associated apathy.

Although the associative structures of both instrumental learning and Pavlovian conditioning underlie goal-directed behaviors (Rescorla, 1987), voluntary wheel running is driven primarily by instrumental learning. Instrumental learning is characterized by the development of an association between a stimulus and response (for review, Colwill & Rescorla, 1986). Early instrumental learning studies (Thorndike, 1898; Thorndike, 1911) led to the development of the “Law of Effect”, which states that a stimulus-response association is bolstered when followed by a reward and weakened when followed by an adverse consequence. With specific regards to voluntary wheel running, rodents develop an association between the running wheel (i.e., stimulus) and successful wheel revolution (i.e., response). The pronounced escalation of daily running during the acquisition phase illustrates the strengthening of the stimulus-response association, independent of genotype, in response to the naturally rewarding properties of running (e.g., Kagan & Berkun, 1954; Collier & Hirsch, 1971; Greenwood et al., 2011; Basso & Morrell, 2015). The maintenance phase of voluntary wheel running is consistent with another key feature of the “Law of Effect”: an organism is compelled to make a response whenever the stimulus is present (Thorndike, 1898; Thorndike, 1911). The systematic manipulation of multiple variables, including time-of-day and wheel access, in combination with a bout microstructural analysis afforded an opportunity to delineate motivational alterations resulting from HIV-1 viral proteins.

Chronic HIV-1 viral protein exposure induced prominent apathetic behaviors for a natural reward, indexed using voluntary wheel running. At the macrostructural level, apathetic behaviors in the HIV-1 Tg rat were characterized by time-of-day dependent alterations in the development of voluntary wheel running behavior. Of specific note is the decreased number of meters run per session in HIV-1 Tg animals during the maintenance phase of nocturnal voluntary wheel running; an observation that supports a decreased reinforcing efficacy of voluntary wheel running in HIV-1 rats. Notably, the HIV-1 Tg rat also exhibited a diminished reinforcing efficacy of sucrose (Bertrand et al., 2018; McLaurin et al., 2021b) supporting a generalizable decrease in motivation for natural rewards. At the microstructural level, HIV-1 Tg animals exhibited a decreased number of running bouts relative to controls supporting a profound alteration in the self-initiation of spontaneous behaviors, one of the key steps for goal-directed behaviors.

Given the multidimensional nature of apathy (Marin, 1991a), investigating individual dimensions (e.g., intention, initiation or execution, and evaluation; initiation, planning, and motivation) is fundamental to our understanding of HIV-1 associated apathy. The evaluation of apathy in HIV-1 seropositive individuals (e.g., Tate et al., 2003; Paul et al., 2005; Kamat et al., 2012; Kamat et al., 2016) has primarily relied upon the classic Apathy Evaluation Scale (AES; Marin et al., 1991b) and the Frontal Systems Behavior Scale (FrSBe; formerly known as the Frontal Lobe Personality Scale; Grace et al., 1999). The importance and utility of these assessments is undeniable, as both scales exhibit high reliability and validity (for review, Clarke et al., 2011); they are limited, however, by their measurement of apathy as a unidimensional construct. Specifically, although the AES includes items evaluating the affective, behavioral, and cognitive domains of apathy, utilization of a composite score (Marin, 1991b) precludes examination of the unique dimensions of apathy. The future implementation of additional scales designed to evaluate the multidimensional nature of apathy (e.g., Apathy Inventory, Robert et al., 2002; Dimensional Apathy Scale, Radakovic & Abrahams, 2014) may further enhance our understanding of HIV-1 associated apathy.

An examination of potential neural mechanisms underlying HIV-1 associated apathy, including synaptic dysfunction in MSNs of the NAc, also addresses a key knowledge gap. MSNs of the NAc are characterized by high densities of dendritic spines, which serve as the main postsynaptic compartment of excitatory synapses (Harris & Kater, 1994). Dendritic spines located on proximal dendrites of MSNs primarily receive collaterals from other MSNs, while those on distal dendrites are innervated by glutamatergic projections from the PFC and dopaminergic afferents from the VTA (Spiga et al., 2014). Furthermore, glutamatergic afferents are targeted at the dendritic spine head (Smith et al., 1994), whereas the dendritic spine neck receives dopaminergic projections (Freund et al., 1984). Additionally, the morphological parameters of dendritic spines may afford critical insight into synaptic function (Harris & Stevens, 1988; Harris & Stevens, 1989; Arellano et al., 2007).

Following a history of voluntary wheel running, HIV-1 Tg animals exhibited prominent synaptic dysfunction, evidenced by alterations in neuronal and dendritic spine morphology in MSNs of the NAc. The leftward distributional shift, whereby HIV-1 Tg animals exhibited an increased prevalence of dendritic spines on more proximal dendrites relative to control animals, supports a prominent alteration in synaptic connectivity. HIV-1 Tg animals also exhibited an immature (i.e., ‘stubby’) dendritic spine phenotype, evidenced by a population shift towards increased dendritic spine head and neck diameter, relative to control animals. Both the preponderance of dendritic spines on more proximal branches and the shift towards a ‘stubby’ dendritic spine phenotype (characterized by the absence of a dendritic spine neck; Peters & Kaiserman-Abramof, 1970) support prominent alterations in dopaminergic neurotransmission. Collectively, the decreased dopamine levels commonly observed in HIV-1 seropositive individuals (e.g., Kumar et al., 2009; Kumar et al., 2011) and biological systems utilized to model HAND (e.g., Jenuwein et al., 2004; Ferris et al., 2009; Denton et al., 2021) may result from and/or lead to alterations in dendritic spines (McLaurin et al., 2021a).

Utilization of multiple regression analyses afforded a critical opportunity to systematically establish synaptic dysfunction in the NAc as a neural mechanism underlying the self-initiation of spontaneous behaviors. Multiple regression analyses evaluate the relationship between one dependent variable and multiple independent variables (Tabachnik & Fidell, 2014). Measures of synaptic dysfunction explained 42%, 68.5% and 54.5% of the variance in the number of running bouts (i.e., self-initiation of spontaneous behaviors) during the diurnal, nocturnal, and deprivation/reinstatement phases, respectively. Neuroimaging studies in HIV-1 seropositive individuals corroborate the importance of the NAc as a neural mechanism underlying HIV-1 associated apathy, whereby the volume of the NAc explained 34.8% of the variance in selfreported apathetic behaviors (Paul et al., 2005). Apathy in HIV-1 seropositive individuals has also been associated with white matter abnormalities in brain regions associated with the fronto-basal ganglia circuit (Kamat et al., 2014). Critically, the importance of the NAc in apathy does not preclude the involvement of other brain regions, specifically those of the fronto-striatal circuit.

One of the notable advantages of the HIV-1 Tg rat is its utility for investigating the debilitating neurocognitive impairments (e.g., Vigorito et al., 2007; Moran et al., 2014; Repunte Canonigo et al., 2014; McLaurin et al., 2019) and affective (e.g., Nemeth et al., 2014; Bertrand et al., 2018; Denton et al., 2021; McLaurin et al., 2021b) alterations associated with HIV-1. The HIV-1 Tg rat, developed by Reid et al. (2001), constitutively expresses seven of the nine HIV-1 viral proteins (the *gag* and *pol* genes are deleted) under the control of the natural HIV-1 promoter. No significant health disparities (i.e., similar growth rates, intact gross-motoric system function) were observed in HIV-1 Tg rats, consistent with prior reports (e.g., Somatic Growth: Peng et al., 2010; Moran et al., 2013; Gross-Motoric System Function: McLaurin et al., 2018b). The present study also provided strong evidence for the integrity of the circadian rhythm in the HIV-1 Tg rat. Specifically, when HIV-1 Tg and control animals were evaluated in the diurnal return phase at the end of experimentation, the average running distance was statistically indistinguishable from the diurnal phase (**Supplementary Figure 2**); observations which support the ability of the HIV-1 Tg rat to adjust to time-of-day manipulations. The prominent apathetic behaviors in the HIV-1 Tg rat, therefore, were not confounded by adverse clinical manifestations.

Taken together, chronic HIV-1 viral protein exposure induces prominent apathetic behaviors characterized by alterations in the self-initiation of spontaneous behaviors. Profound synaptic dysfunction, evidenced by alterations in synaptic connectivity and a shift towards an immature dendritic spine phenotype, mechanistically underlies the self-initiation of spontaneous behavior. Critically, the establishment of a fundamental neural mechanism underlying apathy affords a key target for the development of novel therapeutics and cure strategies for affective alterations associated with HIV-1.

## ACKNOWLEDGEMENTS

This work was supported in part by grants from NIH (National Institute on Drug Abuse, DA013137; National Institute of Child Health and Human Development HD043680; National Institute of Mental Health, MH106392; National Institute of Neurological Disorders and Stroke, NS100624) and the interdisciplinary research training program supported by the University of South Carolina Behavioral-Biomedical Interface Program.

## AUTHOR CONTRIBUTIONS

MNC conducted the behavioral testing of the animals, neuroanatomical assessments and preliminary data analysis, KAM conducted final statistical analyses, graphed the data and wrote the manuscript, HL conducted neuroanatomical assessments, SBH edited the manuscript, and RMB and CFM oversaw the conduct of the research and contributed to all aspects of the project as needed for its completion. All authors read and approved the final manuscript. MNC is now a postdoctoral fellow at the Uniformed Services University of the Health Sciences, Department of Pathology, Bethesda, Maryland.

