## Supplementary Figures 1 and 2 for "SYNAPTIC DYSFUNCTION UNDERLIES ALTERATIONS IN THE INITIATION OF GOAL-DIRECTED BEHAVIORS: IMPLICATIONS FOR HIV-1 ASSOCIATED APATHY"

Program in Behavioral Neuroscience

Department of Psychology

University of South Carolina

Columbia, SC 29208

*These authors contributed equally.

**Address proofs and correspondence to:**

Rosemarie M. Booze, Ph.D.

Carolina Trustees Professor and Bicentennial Endowed Chair of Behavioral Neuroscience

Department of Psychology

1512 Pendleton Street

University of South Carolina

Columbia, SC 29208

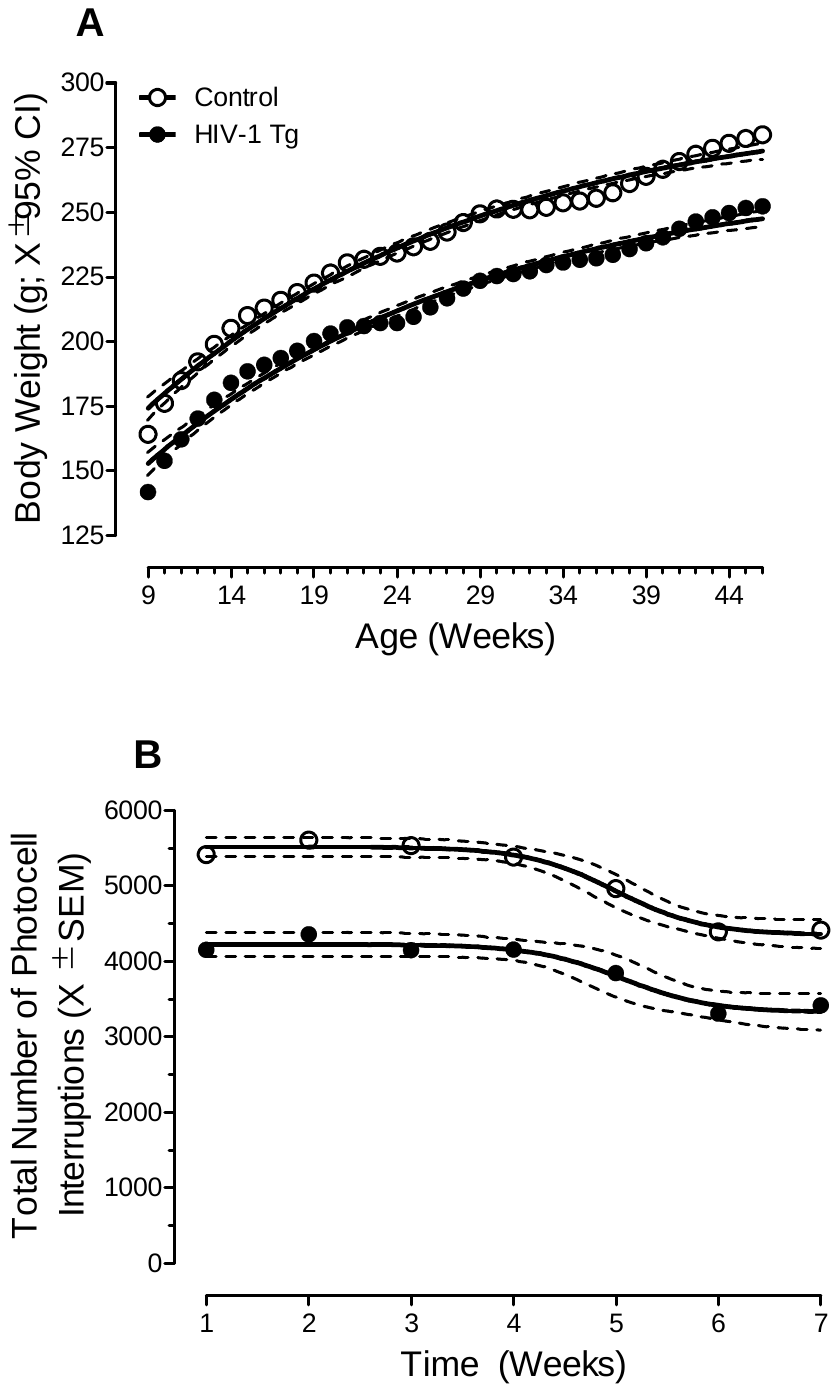

**Supplementary Figure 1.** Experimental controls, including measures of somatic growth (**A**) and gross-motoric system function (**B**), are illustrated as a function of genotype (HIV-1 Tg vs. Control; ±95% Confidence Intervals). With regards to somatic growth, HIV-1 Tg animals weighed significantly less than control animals throughout the duration of the experiment; albeit no significant differences in the rate of growth were observed (**A**). The integrity of gross-motoric system function was evidenced via the examination of locomotor activity, whereby the number of photocell interruptions for both HIV-1 Tg and control animals were greater than zero across all testing sessions.

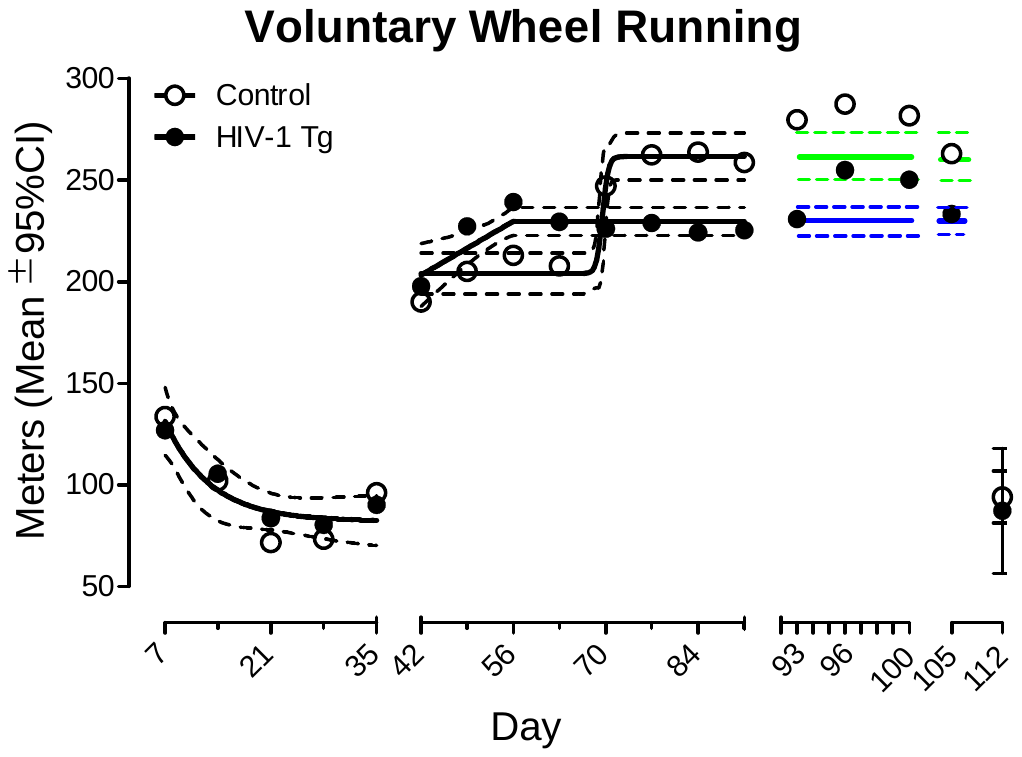

**Supplementary Figure 2.** Voluntary wheel running throughout the experiment is illustrated as a function of genotype (HIV-1 Tg vs. Control; ±95% Confidence Intervals or SEM).The total number of meters run during an individual daily session were averaged across seven days. Breaks along the x-axis indicate different phases of the experiment (Diurnal Phase: Days 7-35; Nocturnal Phase: Days 42-91; Deprivation/Reinstatement Phase: Days 92-100; Nocturnal Phase: Days 101-105; Diurnal Return: Days 106-112). The average running distance during the Diurnal Return was statistically indistinguishable from the diurnal phase, supporting the integrity of the circadian rhythm in the HIV-1 Tg rat. The green (Control) and blue (HIV-1 Tg) dashed lines represent the extension of the 95% confidence intervals from the nocturnal phase.
